# Structural and evolutionary insights into DAF-12 interactions with transcriptional coactivators in parasitic nematodes

**DOI:** 10.64898/2026.03.11.711005

**Authors:** Manon Mallet, Yéléna Martin, João E. Carvalho, Eva Guchen, Remy Betous, Cherine Bechara, Anne Lespine, Michael Schubert, William Bourguet, Albane le Maire

## Abstract

Parasitic nematodes infect billions of humans and livestock worldwide, causing major health and economic burdens, while the spread of anthelmintic resistance threatens current control strategies. A critical step in parasite infection is the resumption of development of infective third-stage larvae (iL3) upon host entry, a process controlled by the nuclear receptor DAF-12. Activation of DAF-12 by dafachronic acids promotes developmental progression and reproductive maturation, making this receptor an attractive therapeutic target. However, the molecular mechanisms governing DAF-12 activation, particularly transcriptional coactivator recruitment, remain poorly understood.

Here, we combined biophysical, cellular, structural, and bioinformatic approaches to investigate coactivator recognition by DAF-12 from the parasitic nematodes *Brugia malayi* and *Haemonchus contortus*. Crystal structures of ligand-bound DAF-12 ligand-binding domains in complex with coactivator-derived peptides reveal conserved features of ligand-dependent coactivator recruitment shared with mammalian nuclear receptors. In addition, we uncover previously unrecognized interaction features, including motif-specific contacts that extend beyond the canonical LXXLL binding mode of coactivators and distinct patterns of DAF-12 conservation across nematode clades. Structure-guided analyses redefine the interaction motif of the only described parasite-specific coactivator DIP-1 and suggest novel candidate motifs for DAF-12-interacting proteins.

Together, these findings establish the structural basis of coactivator binding to nematode DAF-12 and provide mechanistic insight into the transcriptional regulation underlying parasite development. These results expand current understanding of nuclear receptor signaling in parasitic nematodes and provide a framework for the future design of strategies aimed at disrupting DAF-12 activation as a potential antiparasitic approach.

**Author Summary:** Parasitic nematodes infect billions of people and livestock worldwide, causing major health and economic burdens, while increasing resistance threatens current treatments. These parasites rely on a developmental switch that allows infectious larvae to resume growth inside their host, a process controlled by the nuclear receptor DAF-12. Blocking this pathway could prevent parasites from establishing infection. However, the molecular mechanisms regulating DAF-12 activation remain poorly understood. Here, we investigate how DAF-12 from two parasitic nematodes, *Brugia malayi* and *Haemonchus contortus*, interacts with transcriptional coactivators that enable gene activation, using a combination of biophysical, cellular, structural, and bioinformatic approaches. We identified conserved features of ligand-dependent coactivator recruitment shared with mammalian nuclear receptors as well as nematode-specific interaction mechanisms that vary across evolutionary clades. These findings provide new insights into the structural basis of coactivator binding to DAF-12 and advance our understanding of a key pathway controlling parasitic nematode development.

## Introduction

A vast number of nematode parasites are known to infect humans, causing an important public health problem. Parasitic nematodes infect nearly three billion people worldwide, and are responsible for several neglected tropical diseases (NTDs) prioritized by the World Health Organization [1], among them river blindness, cutaneous and lymphatic filariasis caused by filarial worms such as *Onchocerca volvulus* or *Brugia malayi*. Beyond their impact on human health, parasitic nematodes are also major pathogens in domesticated animals, as seen with *Dirofilaria immitis* and *Dirofilaria repens*, which cause zoonotic dirofilariasis. Additionally, parasitic nematodes severely alter animal welfare and food production, leading to substantial economic loss for the livestock industry. The global veterinary market for anthelmintic pharmaceuticals exceeds $3.5 billion annually, further exacerbated by mortality and productivity loss. Among the most problematic species, the gastrointestinal nematode *Haemonchus contortus*, infecting small ruminants, is the most prevalent species due to its high pathogenicity, prolificacy, and genetic variability. These traits increase disease burden and accelerate the development of resistance to multiple anthelmintic treatments, posing a critical threat to sustainable livestock management [2].

Parasitic nematodes belong to distinct evolutionary clades, with markedly different and complex life cycles involving multiple developmental stages and host interactions. For instance, *B. malayi* belongs to the clade III superfamily *Filarioideae* and exhibits a complex life cycle involving multiple developmental stages in both an arthropod vector and the definitive mammalian host. *B. malayi* infects the human lymphatic system, where adult females release microfilariae (L1 larvae) into the bloodstream. They are ingested by mosquitoes, in which the microfilariae develop into infective L3 larvae (iL3), and re-enter the human host upon a mosquito bite, maturing into adults in the lymphatic system. By contrast, *H. contortus*, a soil-borne nematode belonging to the clade V superfamily *Trichostrongyloidea*, follows a free-living phase before ending its life cycle within a ruminant host. Adults localize in the abomasum, where they shed eggs in the feces. These eggs hatch into L1 larvae and develop into iL3 larvae, which mature into adults after ingestion by a new host. Despite these differences in life cycles, parasitic nematodes share common features: (i) they enter their mammalian host as developmentally arrested iL3 larvae, which resume development upon infection, and (ii) subsequently undergo molts to adapt to the host environment and mature into reproductive adults. The iL3 stage is functionally analogous to the dauer larval stage of the free-living nematode *Caenorhabditis elegans*, characterized by larval developmental arrest and resistance to harsh environments.

In *C. elegans*, DAF-12 plays a key role in regulating entry into and exit from the quiescent dauer stage [3]. This process is controlled by dafachronic acids (DAs), which act as molecular signals to trigger developmental transitions. Under unfavorable conditions, such as food deprivation, *C. elegans* larvae stop producing DAs and enter into quiescence. When environmental conditions improve, DA synthesis resumes, activating DAF-12, which in turn induces expression of reproductive development genes and facilitates dauer exit. Consequently, inhibiting *C. elegans* DAF-12 activation prevents exit from quiescence and stops reproductive development [4]. Similarly, a large body of evidence supports the critical role of DAF-12 in regulating the infectious processes of various parasitic nematodes [5–11], by controlling the transition from the L3 to the adult stage. *B. malayi* and *D. immitis* DAF-12 bind the natural Δ4-dafachronic acid (Δ4-DA) with significantly higher affinity than their counterparts in non-filarial nematodes, such as *H. contortus* and *C. elegans* [12]. This ligand triggers the exit from the iL3 to the adult stage upon infection of the mammalian host. Thus, inhibiting DAF-12 in these nematodes prevents iL3 larvae from resuming development, thereby blocking the parasites from settling in their host and interrupting growth and reproduction of the worms. Targeting DAF-12 thus represents a valid option for the development of novel therapeutic strategies to combat nematode parasites. Importantly, the potential therapeutic benefit of targeting DAF-12 has already been validated in *Strongyloides stercoralis* [13].

DAF-12 is a ligand-regulated transcription factor belonging to the nuclear receptor superfamily, whose activity is tuned by transcriptional coregulators (coactivators and corepressors). While many coregulators have been identified and found to be associated with numerous diseases in humans [14], only a few have been described in nematodes. A parasite-specific coactivator, called DIP-1, required for DAF-12 ligand-dependent transcriptional activity has been identified in *S. stercoralis* [9], whereas DIN-1, a corepressor that binds unliganded DAF-12 to inhibit programs of reproductive development, diapause, fat storage, and long life in unfavorable conditions has been found in *C. elegans* [15]. The hallmark of nuclear receptors is the presence of a conserved DNA binding domain (DBD) which binds to specific promoters of target genes and a ligand binding domain (LBD) that binds small ligands. Ligand-activated nuclear receptors undergo conformational changes, primarily involving the reorientation of the LBD C-terminal activation helix H12, which facilitates the recruitment of transcriptional coactivators at a conserved interaction surface, called activation function 2 (AF-2), to modulate the transcription of target genes.

No antagonists that can prevent coactivator recruitment to DAF-12 and subsequently its activation have been identified to date. In particular, the development of antagonists based on dafachronic acids, that are the ligands of DAF-12 in different nematodes [11–13] poses substantial practical challenges. The structural complexity of these sterol-derived molecules renders their chemical synthesis and diversification technically demanding and economically costly, thereby limiting their utility as starting points for drug development efforts. Additionally, structural insights on DAF-12 proteins remain limited to LBD characterizations from two parasitic nematode species [10,16], and the repertoire of nematode coactivators is largely undefined. This incomplete understanding of the molecular mechanisms governing DAF-12 activation and regulation explains why progress on novel therapeutic strategies targeting this receptor has been limited.

To better elucidate the mechanism of transcriptional coactivator recruitment by nematode DAF-12, we combined biophysical, cellular, and structural analyses of *B. malayi* and *H. contortus* DAF-12 LBDs together with a comprehensive bioinformatic analysis of DAF-12 sequences across nematodes. We first identified conserved features of ligand-dependent coactivator recruitment shared with mammalian nuclear receptors, which were validated by the determination of the crystal structures of *B. malayi* and *H. contortus* DAF-12 LBDs in complex with mammalian coactivator-derived peptides. Moreover, we uncovered DAF-12-specific interactions with coactivators, displaying varying degrees of conservation across nematode clades. In addition to redefining the interaction motif of DIP-1, we propose novel motifs of coactivators that may interact with DAF-12, providing a framework for the identification of transcriptional coactivators in nematodes. Together, our findings, in complement to the studies on *Ancylostoma ceylanicum* and *S. stercolaris* DAF-12 [10,16], establish the structural basis of coactivator binding to DAF-12, advancing our understanding of the molecular mechanisms of action of this key regulator of nematode development.

## Results

### *B. malayi* and *H. contortus* DAF-12 LBD purification and ligand binding

We structurally and functionally characterized the LBD of DAF-12 from a filarial parasite, *B. malayi* (*Bma*) and a non-filarial parasite, *H. contortus* (*Hco*), which share 55% sequence identity [12] (Fig 1A). Both proteins were expressed in a bacterial system and purified to high purity (S1A and B Fig). Initial attempts to express the *Hco*DAF-12 LBD yielded a low amount of soluble protein. To improve solubility, we introduced four cysteine-to-serine mutations (C538S, C592S, C610S, C646S), following an approach previously used for *A. ceylanicum (Ace*) DAF-12 [16] (S1C Fig). Due to their location in the LBD out of the ligand and coregulator binding sites, these mutations are not expected to affect ligand or coregulator binding. The mutated variant markedly increased *Hco*DAF-12 solubility, yielding over 5 mg of pure protein per liter of culture and was thus used for all the following biophysical and structural analyses. The structural integrity of the purified proteins was verified by circular dichroism (CD) and dynamic light scattering (DLS). CD spectra indicated that they have a high 𝛼-helical content (60.1% for *Bma*DAF-12 and 74.1% for *Hco*DAF-12) (S1D Fig), consistent with the folded conformation of a nuclear receptor LBD. DLS measurements showed a hydrodynamic radius of 2.4 and 2.6 nm for *Bma*DAF-12 and *Hco*DAF-12, respectively (S1E and F Fig), consistent with a monomeric form of the DAF-12 LBDs in solution.

**Figure 1.**
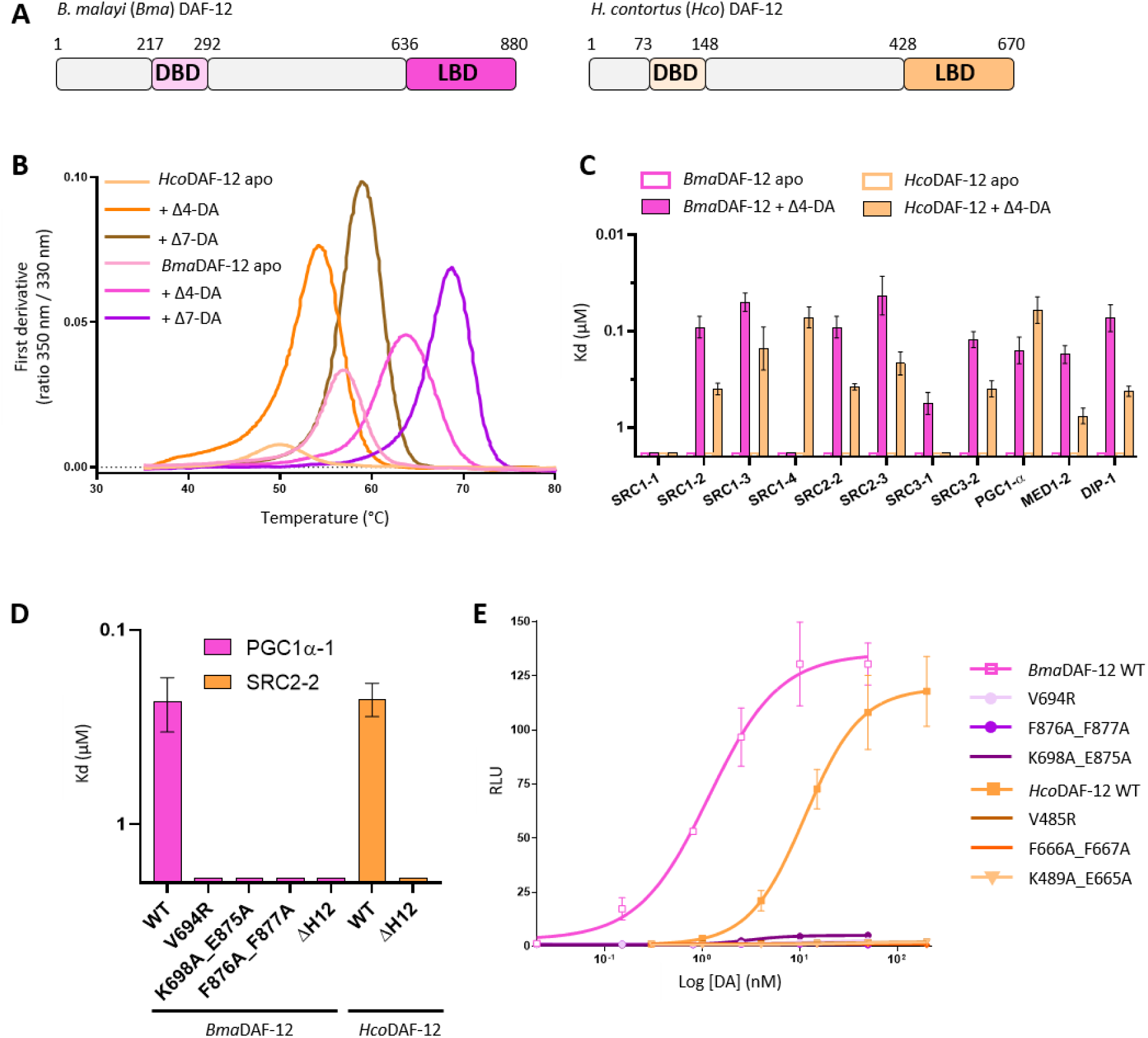
Ligand and coactivator binding properties of *Brugia malayi* (*Bma*) and *Haemonchus contortus* (*Hco*) DAF-12 ligand binding domains. **A.** Schematic representation of *Bma*DAF-12 and *Hco*DAF-12 showing the DNA binding domain (DBD) and ligand binding domain (LBD) limits. **B.** Nano-DSF analysis of ligand binding to *Bma*DAF-12 and *Hco*DAF-12 LBDs. Δ4- and Δ7-dafachronic acids (Δ4-DA and Δ7-DA) or DMSO (apo conditions) were added in one molar equivalent to the proteins. The first derivative of the ratio between intrinsic fluorescence at 350 nm and 300 nm is plotted against the temperature. The experiments were repeated three times independently. **C.** Affinities of *Bma*DAF-12 and *Hco*DAF-12 LBDs for different peptides derived from coactivators, in the absence and in the presence of Δ4-DA. Dissociation constants (Kd in µM) were derived from fluorescence anisotropy experiments. **D.** Affinities of wild-type (WT) and mutant *Bma*DAF-12 for the PGC1α-1 peptide (pink), and of *Hco*DAF-12 and *Hco*DAF-12ΔH12 for the SRC2-2 peptide (orange). Dissociation constants (Kd in µM) were derived from fluorescence anisotropy experiments. **E.** Dose-response curves for Δ4-DA on the activation of WT and mutant *Bma*DAF-12 (pink) and for Δ7-DA on the activation of WT and mutant *Hco*DAF-12 (orange). NIH3T3 cells were co-transfected with Gal4-DAF-12_LBD and the luciferase gene reporter construct. Transfected cells were incubated with increasing concentrations of Δ4-DA or Δ7-DA. The transactivation activity was assessed by measuring luciferase activity, which was normalized by Renilla luciferase activity for transfection efficiency and expressed as relative light units (RLU). Data representing normalized luciferase activities are plotted with nonlinear regression fit using sigmoidal dose response with variable slope (Prism 6.0, Graph Pad Software, Inc.). Values are means from two independent experiments performed in technical triplicates.

To demonstrate ligand binding to purified *Bma*DAF-12 and *Hco*DAF-12 LBDs, we analyzed their thermal stability, without and with one or three molar equivalents of Δ4-dafachronic acid (Δ4-DA) and (25*S*)-Δ7-dafachronic acid (Δ7-DA), using nano-DSF (Fig 1B, S2A and S2B Fig and S1 Table). Both dafachronic acids significantly stabilized the *Bma*DAF-12 and *Hco*DAF-12 LBDs, even at a 1:1 ligand-to-protein ratio, in agreement with their high potency to active *Bma*DAF-12 and *Hco*DAF-12 [12]. Finally, native mass spectrometry (nMS) analysis of *Bma*DAF-12 incubated with two and a half molar excess of Δ4-DA confirmed complex formation and showed a 1:1 stoichiometry, as evidenced by the appearance of a new distribution corresponding to the expected mass of the protein-ligand complex (29170 Da) (S2C and S2D Fig). Thus, our results confirmed the physical binding of dafachronic acids to *B. malayi* and *H. contortus* DAF-12, in agreement with their functional ligand-dependent activation [12].

### *B. malayi* and *H. contortus* DAF-12 recruit coactivators in a ligand-dependent manner

To analyze the ability of *Bma*DAF-12 and *Hco*DAF-12 LBDs to bind transcriptional coactivators in a ligand-dependent manner, we screened a panel of eleven peptides derived from coactivators using steady-state fluorescence anisotropy experiments. Since transcriptional coregulators from *B. malayi* and *H. contortus* remain uncharacterized, this panel included ten LXXLL-containing peptides derived from mammalian coactivators and one FXXLL-containing peptide derived from *S. stercoralis (Sst)* DIP-1 [9], the only coactivator described to date in nematodes (Fig 1C, Table 1 and S3 Fig).

**Table 1.**
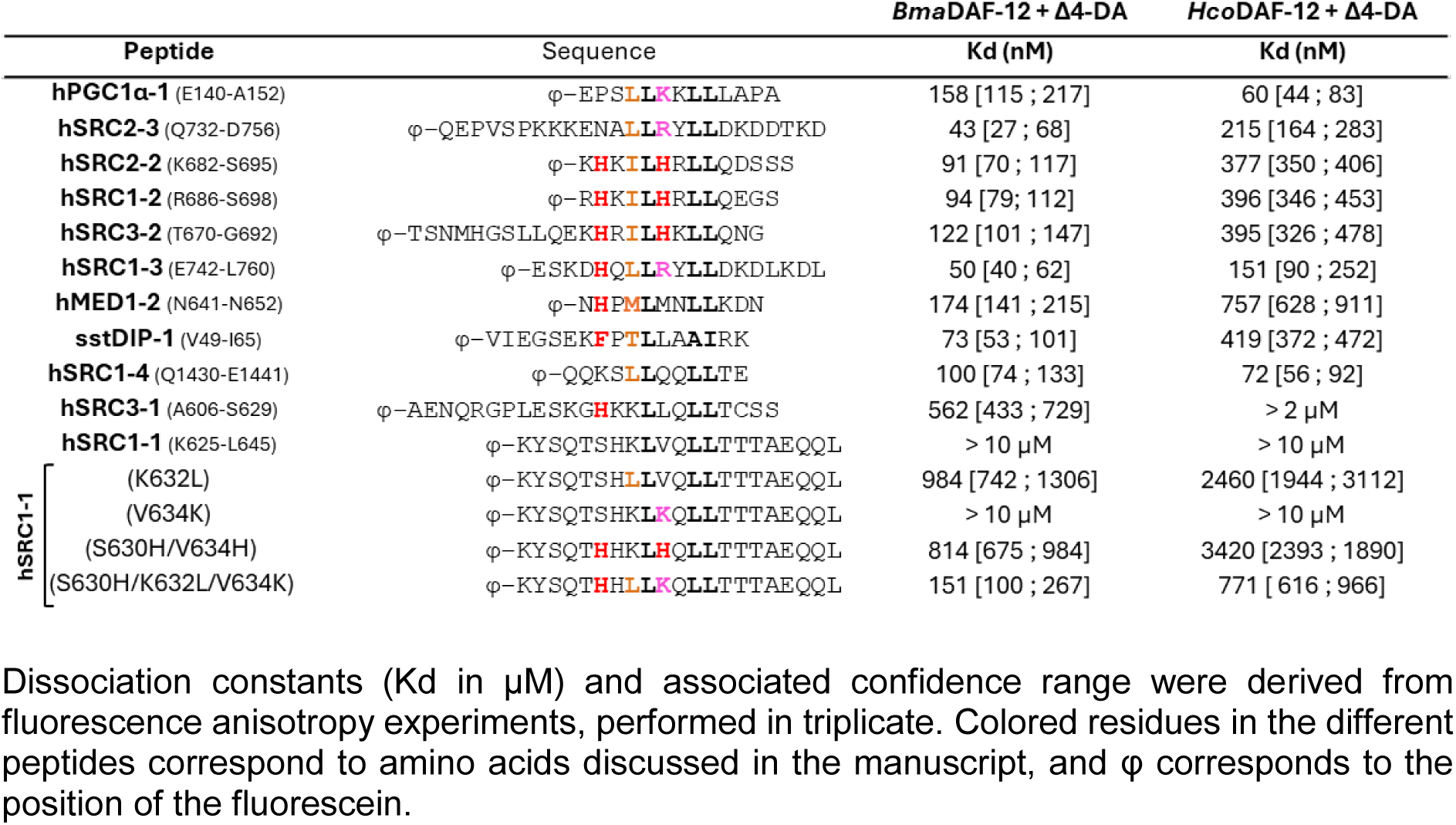
Coactivator binding to *Brugia malayi* (*Bma*) and *Haemonchus contortus* (*Hco*) DAF-12 ligand binding domains, in the presence of Δ4-DA.

As expected, no coactivator binding was measured in the absence of DAs, while in the presence of Δ4-DA, several coactivator-derived peptides were efficiently recruited by *Bma*DAF-12 and *Hco*DAF-12 LBDs. The *Hco*DAF-12 LBD displayed the highest affinities for PGC1⍺-1 and SRC1-4 peptides, as also observed for DAF-12 from *A. ceylanicum*, *Ancylostoma caninum*, *Necator americanus*, and *S. stercolaris* [16]. The *Bma*DAF-12 LBD showed higher affinities for coactivator-derived peptides in general and distinct preferences with affinities ranging from 43 to 174 nM for most of the peptides derived from coactivators belonging to different families. These observations suggest species-specific coactivator preferences among DAF-12 orthologs and suggest that residues flanking the LXXLL motif contribute to both binding affinity and receptor specificity. In addition, both DAF-12 LBDs interacted weakly with the motifs 1 of SRC1 and SCR3. Both *Bma*DAF-12 and *Hco*DAF-12 LBDs also interacted with the *Sst*DIP-1 FXXLL-containing peptide in the presence of Δ4-DA with Kd values of 73 nM and 419 nM, respectively. As expected, Δ7-DA also induced *Sst*DIP-1 binding with comparable affinities (S3C Fig), confirming the agonistic activity of both DAs [12].

These observations also suggest a mode of coactivator binding conserved between mammalian and nematode nuclear receptors. To further substantiate this hypothesis, we performed mutagenesis of DAF-12 conserved residues, which are known to play a critical role in coactivator recruitment by mammalian nuclear receptors [17] (S4A and B Fig). We then performed fluorescence anisotropy assays with *Bma*DAF-12 mutants to assess their binding to the PGC1α-1 peptide (Fig 1D and S3D Fig). Substitution of V694 (V299 in hFXR) within the hydrophobic groove of DAF-12 that interacts with the leucine residues of coactivator peptides with a charged residue (V694R) abolished peptide interaction. Similarly, alanine substitution of the charged clamp residues K698A and E875A (K303 and E467 in hFXR) involved in hydrogen bonds with coactivator peptides eliminated coactivator binding. We next tested whether the active positioning of helix H12 is also stabilized by hydrophobic interactions as observed with I468 and W469 in hFXR. Alanine substitution of the corresponding residues, F876 and F877, prevented the stabilized position of helix H12 and abolished PGC1α-1 binding. To further assess the role of H12, we generated deletion constructs lacking this helix (*Bma*DAF-12ΔH12 and *Hco*DAF-12ΔH12). These variants failed to recruit coactivator peptides in the presence of Δ4-DA (Fig 1D, S3D and E Fig). Consistent with these findings, luciferase-based transactivation assays conducted in NIH3T3 cells showed that none of the *Bma*DAF-12 mutants exhibited transcriptional activity in the presence of Δ4-DA, in contrast to robust luciferase expression by wild-type *Bma*DAF-12 (EC50 = 1.2 nM) (Fig 1E). Equivalent mutations in *Hco*DAF-12 (V485R, K489A/E665A, F666A/F667A) (S4 Fig) similarly abrogated receptor activation in response to Δ7-DA, contrary to the strong luciferase expression by wild-type *Hco*DAF-12 (EC50 = 11.3 nM) (Fig 1E). Together, these results underscore the critical role of helix H12 in mediating coactivator recruitment and confirm that DAF-12 orthologs in parasitic nematodes utilize a conserved activation mechanism closely resembling that of mammalian nuclear receptors.

### Crystal structures of *B. malayi* and *H. contortus* DAF-12 LBDs in complex with coactivator peptides

To further confirm the mechanism of ligand and coactivator recruitment by *B. malayi* and *H. contortus* DAF-12, we conducted crystallization trials of various DAF-12/peptide/Δ4-DA complexes and determined the crystal structures of *Bma*DAF-12 and *Hco*DAF-12 LBDs, each in complex with Δ4-DA and a high-affinity mammalian coactivator-derived peptide (PGC1ɑ-1 and SRC2-2, respectively), at resolutions of 1.70 Å and 2.00 Å (Fig 2A-B and S2 Table). Both LBDs adopt the canonical nuclear receptor fold, composed of twelve α-helices arranged in a three-layer sandwich and a β-sheet. Of note, instead of the short loop (four to five residues) typically found between helices H7 and H8 in mammalian nuclear receptors, both DAF-12 LBDs feature an additional short α-helix (H7’). This structural element has also been observed in the *Sst*DAF-12 *and Ace*DAF-12 LBDs [10,16], appearing unique and specific to this nuclear receptor. Structural superposition of the *Bma*DAF-12 and *Hco*DAF-12 LBDs yields a root mean square deviation (RMSD) of 0.946 Å over all Cα atoms, indicating a high degree of structural conservation between the two proteins (S5 Fig). Both structures also exhibit strong similarity to previously determined structures of the *Sst*DAF-12 and *Ace*DAF-12 LBDs (S5 Fig), further supporting the conserved architecture of this receptor across parasitic nematodes.

**Figure 2.**
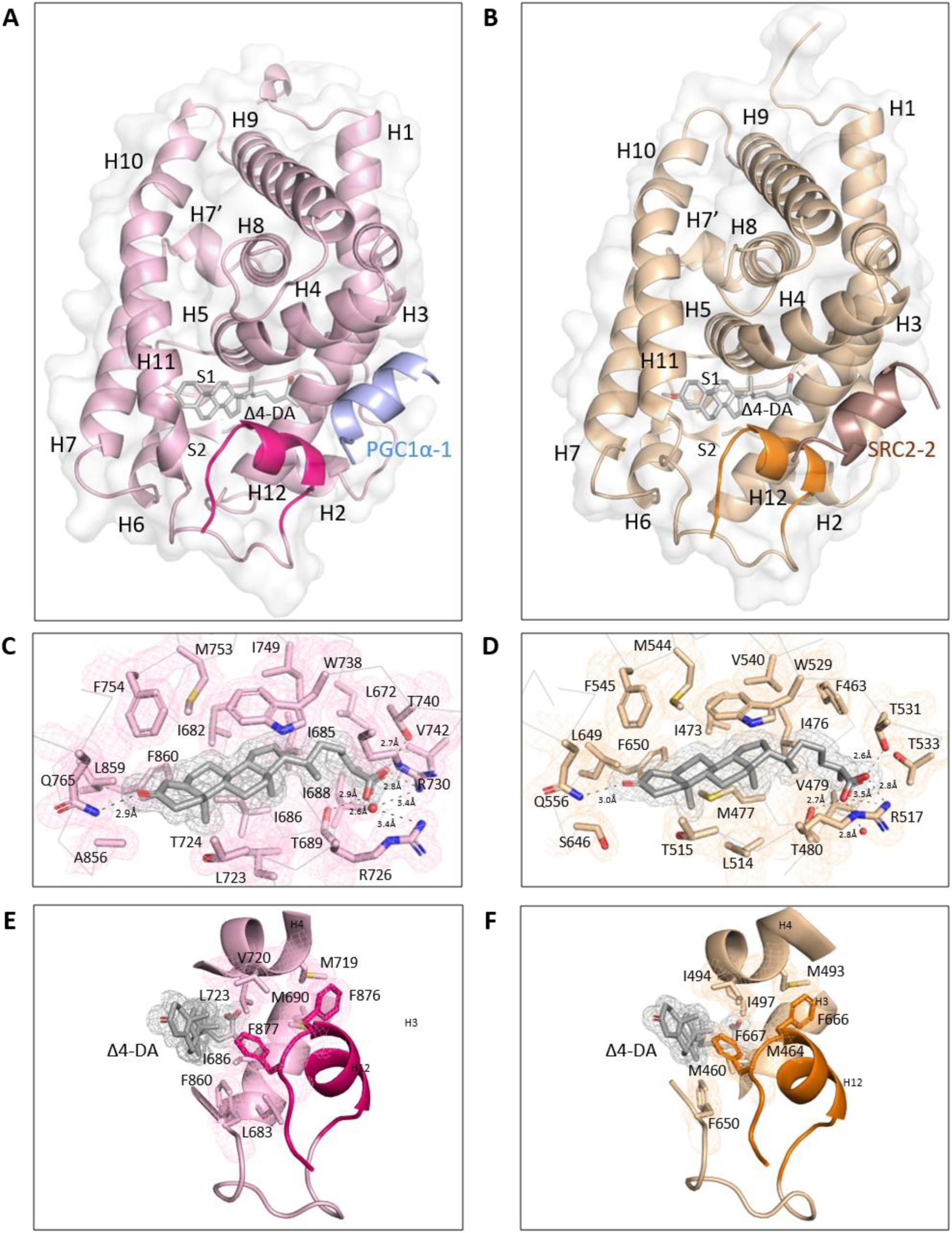
Structural analysis of *Brugia malayi* (*Bma*) *and Haemonchus contortus* (*Hco*) DAF-12 ligand binding domains (LBDs). **A. and B.** Overall structures of *Bma*DAF-12 LBD (light pink) complexed with Δ4-DA (grey) and the coactivator peptide PGC1α-1(blue) (A) and of *Hco*DAF-12 LBD (light orange) complexed with Δ4-DA (grey) and the coactivator peptide SRC2-2 (brown) (B), both in cartoon representation. Helices H12 are highlighted in pink and orange, respectively. **C. and D.** Details of *Bma*DAF-12 (C) and *Hco*DAF-12 (D) ligand binding pockets bound to Δ4-DA. The surrounding amino acids are colored in pink (C) and light orange (D), respectively. The 2*F_0_-F_C_* electron density maps contoured at 1.4 σ are shown for the Δ4-DA and the surrounding amino acids in the corresponding colors. H-bonding of amino acids and water involved in interactions with the C3-keto group and the C27-carboxyl group of Δ4-DA are shown in black dashes and distances are labeled. **E. and F.** Details of helix H12 active conformation stabilization in the *Bma*DAF-12 (E) and *Hco*DAF-12 (F) LBD. Side chains of DAF-12 residues involved in this stabilization are shown as sticks. The 2*F_0_-F_C_* electron density maps contoured at 1.4 σ are shown for the Δ4-DA as well as the stabilizing residues in the corresponding colors.

The ligand binding pocket (LBP) volumes of *Bma*DAF-12 and *Hco*DAF-12 were estimated using KVFinder-web [18] (S3 Table). Consistent with previous observations for other DAF-12 orthologs, the LBPs of around 700 Å³ are relatively small in size and fall within the range observed for human nuclear receptors such as VDR, LXR, and FXR, which are amongst the mammalian nuclear receptors that are most closely related to the DAF-12s. Both DAs have a molecular volume estimated by Chimera X [19] around 420 Å³, showing an occupancy of about 60% of the LBP. The structures of *Bma*DAF-12 and *Hco*DAF-12 LBDs revealed a conserved ligand binding mode, shared with the *Ace*DAF-12 and *Sst*DAF-12 LBDs [10,16] (Fig 2C and D). The C3-ketone group of the ligand forms a hydrogen bond with a highly conserved glutamine residue (Q765 in *Bma*DAF-12 and Q556 in *Hco*DAF-12), similar to what was observed with Q571 and Q637 in *Ace*DAF-12 *and Sst*DAF-12, respectively [10,16]. As previously proposed, the length of the alkyl chain in Δ4-DA is critical for positioning the C27 carboxyl group within the LBP. In *Hco*DAF-12, this carboxyl group forms three H-bonds with T480, T531, and R517 (Fig 2D), whereas, in *Bma*DAF-12, it establishes direct hydrogen bonds with the corresponding threonine residues (T689 and T740) and water-mediated interactions with R726 and R730 (Fig 2C). Additionally, Δ4-DA is stabilized by a network of hydrophobic interactions on both sides of its sterol backbone, involving several conserved hydrophobic residues (Fig 2C and D and S4 Table), further supporting a conserved mechanism of ligand recognition among DAF-12 orthologs.

The ligand Δ4-DA makes direct hydrophobic contacts with F860 in *Bma*DAF-12 and F650 in *Hco*DAF-12, and these phenylalanines in turn create stabilizing π-π stacking interactions with *Bma*DAF-12 F877 and *Hco*DAF-12 F667 in helix H12 (Fig 2E and F). These interactions along with additional hydrophobic contacts between Δ4-DA and surrounding residues within the LBP help stabilize helix H12 in its active conformation, a key structural requirement for coactivator peptide recruitment. Coactivator peptides are anchored to the LBDs primarily through hydrophobic contacts involving the leucine residues of their conserved LxxLL motifs (Fig 3A and B). These leucines engage a hydrophobic groove on the AF-2 surface, formed by helices H3, H4, and H12. Key interacting residues include a valine on helix H3 (V694 in *Bma*DAF-12 and V485 in *Hco*DAF-12), a phenylalanine (F712 in *Bma*DAF-12 and F503 in *Hco*DAF-12) and a leucine (L715 in *Bma*DAF-12 and L506 in *Hco*DAF-12) on helix H4, and a phenylalanine on helix H12 (F876 in *Bma*DAF-12 and F666 in *Hco*DAF-12). In addition, electrostatic interactions occur between the peptide and a characteristic charged clamp formed by a lysine on helix H3 (K698 in *Bma*DAF-12 and K489 in *Hco*DAF-12) and a glutamic acid on helix H12 (E875 in *Bma*DAF-12 and E665 in *Hco*DAF-12) that stabilizes the coactivator complex. These key residues are also conserved in the DAF-12 orthologs [10,16], indicating a broadly conserved coactivator binding mechanism among DAF-12, similar to that of mammalian nuclear receptors.

**Figure 3.**
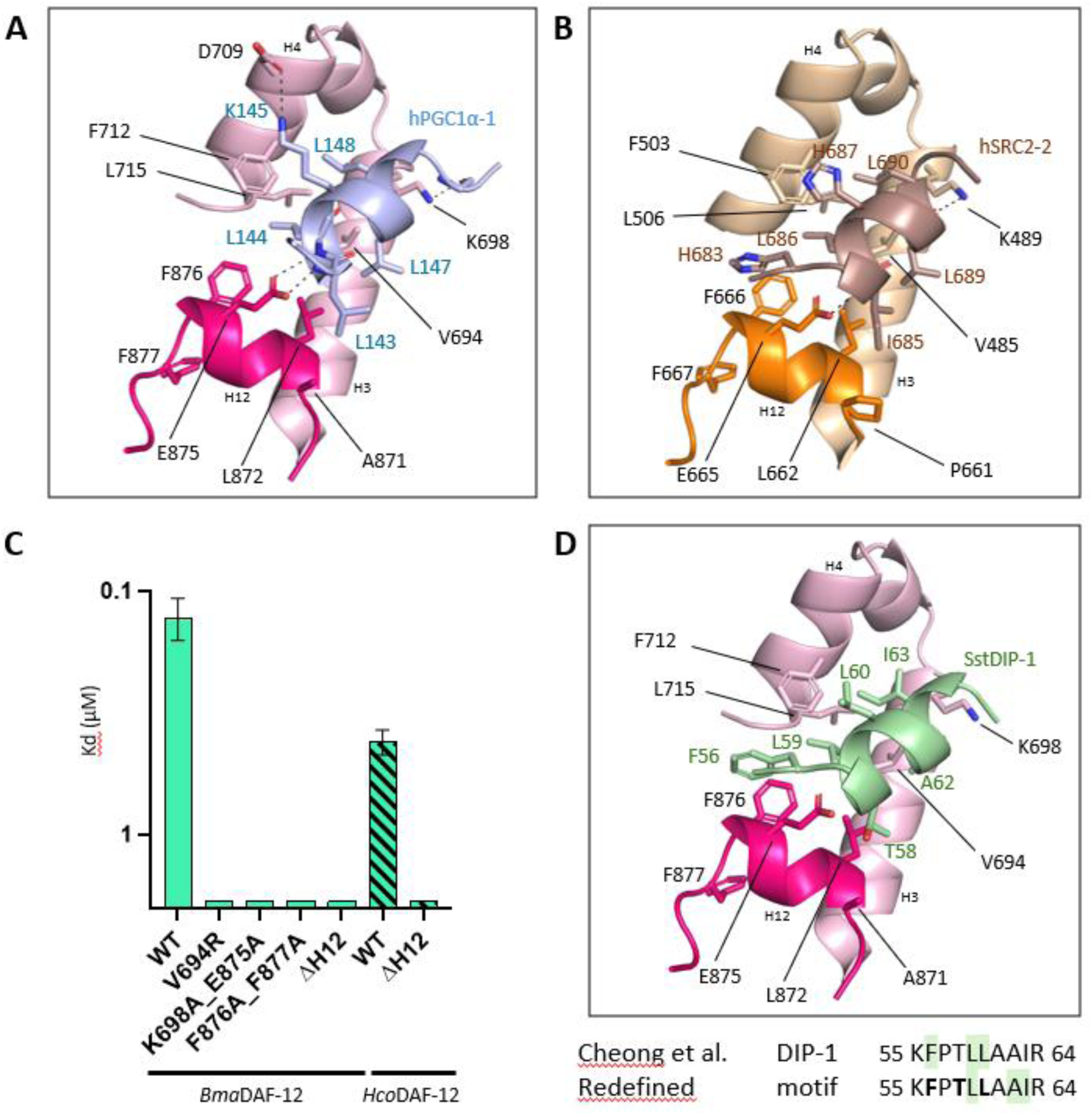
Coactivator binding to the *Brugia malayi* (*Bma*) and *Haemonchus contortus* (*Hco*) DAF-12 ligand binding domains (LBDs). **A. and B.** Close-up view of the *Bma*DAF-12 LBD-PGC1α-1 (A) and *Hco*DAF-12 LBD-SRC2-2 (B) interfaces from the crystal structures of the complexes (presented in this paper). Residues involved in the interaction are shown as sticks and labeled. H-bondings are shown as black dashes. **C.** Affinities of wild-type (WT) and mutant *Bma*DAF-12 (green) and of *Hco*DAF-12 and *Hco*DAF-12ΔH12 (hatched green) for the DIP-1 peptide. Dissociation constants (Kd in µM) were derived from fluorescence anisotropy experiments. **D.** Close-up view of the *Bma*DAF-12 LBD-DIP-1 interface from an AlphaFold model of the complex generated using the Alphafold3 server [20]. Residues involved in the interaction are shown as sticks and labeled. The DIP-1 interaction motif originally defined by Cheong and al. [9] is shown together with the refined motif identified in the present study based on structural analysis. Residues highlighted in green correspond to the minimal interaction motif, whereas residues in bold indicate additional contacts.

### Additional motif-specific interactions between nematode DAF-12 and coactivator peptides

Crystal structures of *Bma*DAF-12 and *Hco*DAF-12 in complex with coactivator-derived peptides reveal motif-specific interactions that extend beyond the canonical LXXLL binding mode. Notably, an electrostatic interaction occurs between lysine K145 in the LKKLL motif of PGC1α-1 and aspartic acid D709 of *Bma*DAF-12 (Fig 3A). A comparable interaction may also contribute to binding of Δ4-DA-bound DAF-12 to SRC1-3 and SRC2-3 peptides (Table 1), which harbor a similar LK/RXLL motif. In the *Hco*DAF-12 complex structure, a π–π stacking is detected between histidine H687 from the LHRLL motif of SRC2-2 and phenylalanine F503 of *Hco*DAF-12, with a distance of about 3.3 Å between the parallel aromatic rings (Fig 3B). This interaction may similarly participate in the recognition of the SRC2-2, SRC1-2, and SRC3-2 peptides by Δ4-DA-bound DAF-12 (Table 1). All motifs interacting with *Bma*DAF-12 and *Hco*DAF-12 also feature a hydrophobic residue immediately preceding the LXXLL motif, yielding a [I/L/M/T]LXXLL consensus (Table 1). This residue (L143 in PGC1α-1 and I685 in SRC2-2) faces hydrophobic residues from helix H12 (alanine A871 and leucine L872 from *Bma*DAF-12 and proline P661 and leucine L662 from *Hco*DAF-12), creating additional hydrophobic contacts and further stabilizing the coactivator peptide on the DAF-12 activation surface (Fig 3A and B). Furthermore, the crystal structure of *Hco*DAF-12 bound to SRC2-2 reveals T-shaped CH–π interactions between the histidine H683, at position −3 relative to the first leucine of the LXXLL motif, and two aromatic residues, F503 and F666, of *Hco*DAF-12 (Fig 3B). Of note, these two phenylalanine residues are conserved in *Bma*DAF-12 (F712 and F876), but not present in mammalian nuclear receptors (S4B Fig). Consistent with these structural observations, the SRC1-1 peptide, which lacks these additional interacting residues, showed no detectable interaction with either DAF-12 receptor.

To evaluate the contribution of individual coactivator peptide residues to DAF-12 binding, we introduced targeted point mutations into the SCR1-1 peptide and quantified binding affinities using fluorescence anisotropy assays (Table 1). Substitution of the lysine at position −1 of the LXXLL motif with a hydrophobic residue (K632L) as well as the double substitution of residues at positions −3 and +1 of the LXXLL motif into histidine (S630H/V634H), resulted in weak but detectable interactions of the corresponding peptides with *Bma*DAF-12 and *Hco*DAF-12 LBDs. In contrast, substitution of the valine within the LVQLL motif by a lysine (V634K) did not induce an interaction of the peptide with the DAF-12 LBDs. Strikingly, combining the three substitutions to generate the S630H/K632L/V634K peptide, which conforms to a HXLLKXLL motif similar to that of the SRC1-3 peptide, yielded high-affinity binding with dissociation constants in the presence of Δ4-DA of approximately 100 nM for *Bma*DAF-12 and 771 nM for *Hco*DAF-12. These mutagenesis experiments thus confirm the critical contribution of residues flanking or within the LXXLL core motif to coactivator binding to DAF-12. On the contrary, these substitutions on SRC1-1 did not induce any interaction with the mammalian nuclear receptors LXRβ and RXRα (S4C Fig), suggesting a specificity towards DAF-12. Collectively, the structural and biochemical analyses uncover novel, motif-specific contacts beyond the canonical LXXLL interaction, providing mechanistic insights into the sequence determinants of DAF-12 coactivator binding in parasitic nematodes.

### A conserved interaction mode between *B. malayi* and *H. contortus* DAF-12 LBDs and *S. stercoralis* DIP-1 and a new definition of its interaction motif

To assess whether the recruitment of *S. stercoralis* DIP-1, the sole coactivator identified in parasitic nematodes, follows a canonical mechanism, we monitored the binding of the DIP-1 peptide spanning residues V49 to K65 to wild-type *Bma*DAF-12 and its mutants (Fig 3C and S3D Fig). Although *Sst*DIP-1 contains a variant motif (FXXLL rather than LXXLL), the same DAF-12 mutations that disrupted binding to the mammalian coactivator peptides (Fig 1D) also abolished interaction with *Sst*DIP-1, supporting the notion that DAF-12 engages both mammalian and nematode coactivators through a conserved structural mechanism. Similarly, the DAF-12 deletion constructs lacking helix H12 (*Bma*DAF-12ΔH12 and *Hco*DAF-12ΔH12) were not able to interact with the DIP-1 peptide, confirming the role of helix H12 in the interaction. To further investigate the binding of *Sst*DIP-1 to DAF-12, we generated AlphaFold [20] models of the *Bma*DAF-12 and *Hco*DAF-12 LBDs in complex with this peptide and compared them to the experimentally determined structures presented in this study (Fig 3D and S6 Fig). Interestingly, this model suggested that the interaction motif of *Sst*DIP-1 is more consistent with a LXXAI motif rather than with the initially proposed FXXLL motif [9]. Leucine L59 and isoleucine I63 play the role of the first and last leucines, respectively, in the classical LXXLL motif by establishing contacts with hydrophobic residues from the AF-2 surface as described above. T58 is also forming hydrophobic contacts with L872 in *Bma*DAF-12 and L662 in *Hco*DAF-12. In addition, the phenylalanine F56 appears to contribute to the interaction in a manner similar to that of histidine H683 in the SRC2-2 peptide, forming a T-shaped CH–π interaction with F712 (H4) and F876 (H12) from *Bma*DAF-12 and with F503 (H4) and F666 (H12) from *Hco*DAF-12. These observations align with the mutagenesis results reported by Cheong and colleagues [9], which identified leucine L59 as a critical determinant for DIP-1 binding, while mutation of the phenylalanine F56 to alanine had a more modest effect on the interaction. To conclude, this detailed structural analysis allowed us to re-define the interaction motif of DIP-1 as a FXXLXXAI motif suggesting that the interaction motifs of nematode coactivators are similar, but not identical, and thus more diverse than their mammalian counterparts.

### DAF-12 LBD is conserved in nematodes from clades III, IV, and V

To determine whether *B. malayi* and *H. contortus* DAF-12 residues implicated in coactivator recruitment are conserved across nematodes, we performed a comprehensive bioinformatic analysis of DAF-12 sequences. We surveyed all nematode genomes available in WormBase [21] and identified DAF-12 orthologs exclusively in species belonging to clades III, IV, and V. A phylogenetic tree based on an alignment of 118 full-length DAF-12 sequences (S7 Fig) closely recapitulated nematode taxonomy, with sequences clustering according to their respective clades. We next examined sequence conservation within the DAF-12 LBDs (Fig 4). A sequence logo analysis revealed a high degree of conservation, particularly in regions corresponding to helices H3, H4, H5, and H12. In contrast, sequence conservation was significantly lower in loop regions. Overall, DAF-12 sequences were most highly conserved among clade III nematodes, whereas greater, though still limited, sequence variability was observed within clade IV and clade V nematodes (S8 Fig).

**Figure 4.**
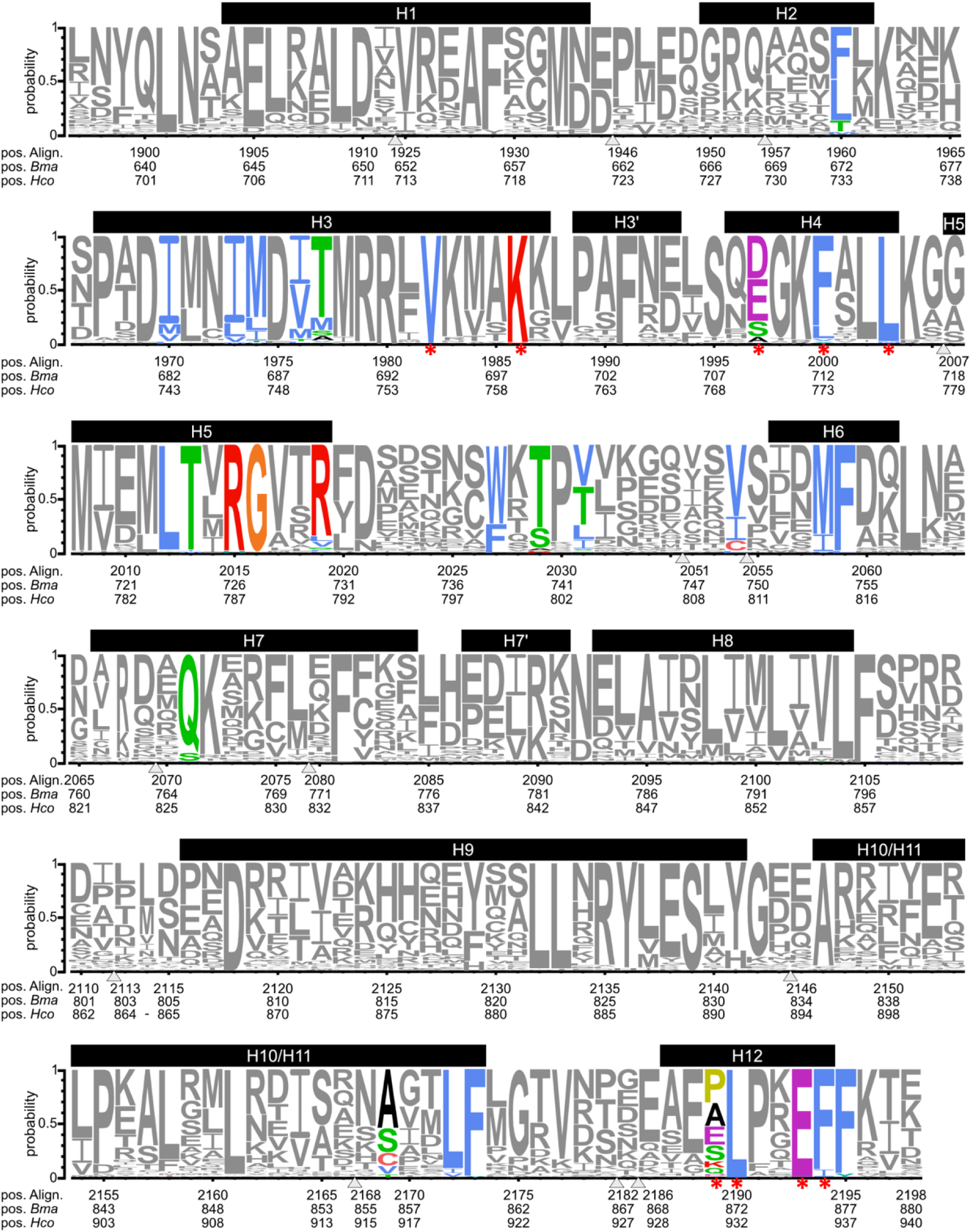
Sequence conservation within DAF-12 ligand-binding domains (LBDs). Sequence logos were generated from positions 1897 to 2198 (corresponding to the LBD) of the untrimmed multiple sequence alignment. Colored residues indicate amino acids experimentally identified as contributing to ligand and coactivator binding, and residues marked by a red asterisk are only involved in coactivator binding. Black horizontal bars refer to secondary structure elements (helices H1 to H12). Arrowheads mark regions that were masked in the alignment (see S7 Fig for the complete alignment).

With respect to residues involved in ligand binding, all the hydrophobic residues lining the LBP were markedly conserved in terms of their hydrophobic character. The arginine corresponding to R726 in *B. malayi* (and R517 in *H. contortus*), which is critical for coordinating the carboxyl group of dafachronic acids, was strictly conserved in all DAF-12 sequences (Fig 4). By comparison, residues that play a more auxiliary role in ligand recognition, such as arginine R730 and threonine T740 (*B. malayi* numbering), displayed some minor variability (S8 Fig). Similarly, the glutamine residue corresponding to Q765 in *B. malayi* (Q556 in *H. contortus*), which contributes to binding of the C3-ketone group of the ligand, was strongly conserved across clades. Together, these observations suggest that DAs are likely to serve as common ligands for DAF-12 across different nematode clades.

Regarding residues involved in coactivator binding, all positions described above to be critical for the recruitment of coactivator-derived peptides were strictly conserved across the DAF-12s from the three clades, with two notable exceptions. First, the aspartic acid corresponding to D709 in *B. malayi*, which forms an electrostatic interaction with K145 of PGC1𝛂-1 (Fig 3A) was highly conserved in clade III nematodes, but showed a notable variability in clades IV and V (S8 Fig). Similarly, alanine A871 in *B. malayi* DAF-12, which faces a hydrophobic residue in the coactivator-derived peptide, was highly conserved in clade III nematodes, but was more variable in clades IV and V, including substitutions by a charged residue (S8 Fig). This suggests that DAF-12 proteins might exhibit clade-specific differences in coactivator preference. Together, these observations on DAF-12 sequences from different nematode clades support a conserved interaction mechanism with coactivators and suggest that residues flanking or within the LXXLL motifs may contribute to the specificity of coactivator binding to DAF-12 and that this specificity might vary between different nematode clades.

## Discussion

In this study, we focused on the structure of DAF-12 receptors from two parasitic nematodes of major biomedical and veterinary importance: *B. malayi*, a filarial nematode causing several NTDs with a huge impact on human health, and *H. contortus*, a gastrointestinal nematode of wild and domesticated ruminants that is highly pathogenic and economically significant. The in-depth structural analysis of DAF-12 from *B. malayi* and *H. contortus* in complexes with transcriptional coactivator-derived peptides combined with mutagenesis experiments and a comprehensive bioinformatic analysis of DAF-12 sequences allowed us to identify key structural features shared across all DAF-12 orthologs. These include the residues lining the ligand binding pocket as well as the canonical charge clamp formed by K698 and E875 in *B. malayi* (K489 and E665 in *H. contortus*), and the hydrophobic groove formed by residues from H3, H4, and H12, both involved in coregulator binding. Comparison with the closest mammalian nuclear receptors, namely VDR, FXR, and LXR, revealed additional DAF-12-specific features within the coactivator binding surface. Notably, the strong conservation of F712 and F876 in *B. malayi* (F503 and F666 in *H. contortus*) suggests nematode-specific interaction determinants that may facilitate selective recruitment of nematode transcriptional coactivators. In addition, we confirmed the agonist activities of Δ4-and Δ7-DAs, two known DAF-12 ligands [3,8,22], in promoting the recruitment of coactivators, in particular that of *S. stercoralis* DIP-1 [9], to *B. malayi* and *H. contortus* DAF-12. Altogether, these results support a conserved mechanism of DAF-12 regulation in parasitic nematodes, in line with the essential role of this nuclear receptor in nematode development [5–11].

With the exception of *Sst*DIP-1, no transcriptional coactivators have been described in nematodes, which limits our ability to fully characterize the regulatory mechanisms governing DAF-12 and other nuclear receptors. Nevertheless, our current findings provide a framework for the identification of DAF-12 coactivators in these species. Specifically, our data allowed us to define a conserved interaction motif in coactivator-derived peptides that interact with DAF-12 and are characterized by the consensus sequence [F/H]X[L/I/T/M]L[K/R/H]X[A/I/L][I/L]. Given that complete genome sequences of a number of different nematodes are available in WormBase [21], this motif provides a basis for a systematic *in silico* screening of potential DAF-12 coactivators. To further refine candidate selection, priority should be given to proteins containing two to three such motifs, in accordance with the architecture of mammalian nuclear receptor coactivators [23].

Previous studies have established that DAF-12 is required in *C. elegans* both to enter into and exit from the dauer diapause, a developmental stage that corresponds to the infective L3 stage in many parasitic nematodes [3,4]. Our findings demonstrate that the mechanism of activation of DAF-12 is highly conserved across parasitic nematodes, underscoring the role of this nuclear receptor in parasite development. This conservation also suggests that targeting DAF-12 signaling represents a promising therapeutic strategy, distinct from current anthelmintic drug targets, for the control of parasitic nematode infections affecting humans, animals, and plants. By elucidating the structural determinants of ligand binding and coactivator recruitment to DAF-12, this study thus provides a framework for the rational design of compounds that selectively target DAF-12 activity, with potential therapeutic applications.

## Materials and Methods

### Ligands and peptides

Δ4-dafachronic acid (Δ4-DA) and (25*S*) Δ7-dafachronic acid (Δ7-DA) were purchased from Cayman and ChemCruz, respectively. The fluorescent peptides labelled with FITC at their N-terminal and listed in Table 1 were purchased from Proteogenix. The unlabelled peptides PGC1𝛼-1 (EEPSLLKKLLLAPA) and SRC2-2 (KHKILHRLLQDSS) used for crystallization were purchased from Ezbiolab.

### Protein purification

DNA sequences corresponding to *Bma*DAF-12 LBD (from L636 to T880, Uniprot entry A0A4E9F2L4), *Hco*DAF-12 LBD (from L428 to E670, Uniprot entry A0A7I4Z2W0) and *Hco*DAF-12 LBD (C538S, C592S, C610S, C646S), *Bma*DAF-12ΔH12 (from L636 to G862), and *Hco*DAF-12ΔH12 (from L428 to G652) have been optimized for *Escherichia coli* expression using the Integrated DNA Technologies (IDT) tool and were purchased at IDT. The sequences were cloned into the pDB-His-TXR-3C vector and expressed using the *E. coli* BL21(DE3) cellular system.

Cells were grown at 37°C in LB medium supplemented with 50 µg/mL kanamycin until OD_600_ reached about 0.6. Expression of T7 polymerase was induced by addition of isopropyl-β-d-thiogalactoside (IPTG) to a final concentration of 0.5 mM for *Bma*DAF-12 LBD and of 1mM for *Hco*DAF-12 LBD. After an additional incubation of 24 hours at 16°C, cell cultures were harvested by centrifugation at 6,000 g for 20 minutes. The cell pellet was resuspended in buffer A (20 mM Tris-HCl pH 7.5, 150 mM NaCl for *Bma*DAF-12 and *Bma*DAF-12ΔH12; 20 mM Tris-HCl pH 7.5, 200 mM NaCl, 0.2% IGEPAL for *Hco*DAF-12 and 0.5% IGEPAL for *Hco*DAF-12ΔH12), supplemented with a protease inhibitor mixture (complete, mini, EDTA-free tablet) and 10 µg/mL lysozyme. The suspension was then lysed by sonication and centrifuged at 40,000 g and 6°C for 30 minutes. The supernatant was loaded onto a 5 mL Ni^2+^-affinity column, preequilibrated with buffer A containing 10 mM imidazole, using the Akta purifier system. The column was washed with 20 column volumes (CV) of buffer A containing 10 mM imidazole and 20 CV of buffer A containing 20 mM imidazole. Bound proteins were incubated overnight at 4°C with an excess of PreScission protease to cleave the His-TRX tag. Unbound protein was eluted with buffer A containing 30 mM imidazole. The protein was further purified using a Superdex 75 16/60 gel filtration column preequilibrated with buffer C (20 mM Tris-HCl pH 7.5, 150 mM NaCl, 5% (v/v) glycerol, 2 mM DTT for *Bma*DAF-12 and *Bma*DAF-12ΔH12; 20 mM Tris-HCl pH 7.5, 150 mM NaCl, 5% (v/v) glycerol, 1 mM DTT for *Hco*DAF-12 and 50 mM Tris pH8, 500 mM NaCl, 5% (v/v) glycerol for *Hco*DAF-12ΔH12). The protein-containing fractions were pooled and concentrated using Amicon-Ultra 10000 MXCO centrifugal filter units (Millipore).

### Nano-differential scanning fluorimetry (Nano-DSF)

Nano-differential scanning fluorimetry (Nano-DSF) experiments were performed on a Tycho NT.6 instrument (NanoTemper) to analyze the thermal stability of the proteins. Protein samples were prepared at 5 µM for *Bma*DAF-12 in SEC buffer (20 mM Tris-HCl pH 7.5, 150 mM NaCl, 5% (v/v) glycerol, and 2 mM DTT) and at 20 µM for *Hco*DAF-12 in SEC buffer (20 mM Tris-HCl pH 7.5, 150 mM NaCl, 5% (v/v) glycerol, and 1 mM DTT). Samples were prepared in the presence of one and three molar equivalents of ligands initially prepared at 20 mM in DMSO. For the reference samples, DMSO alone was added to a final concentration identical to the one in the presence of the ligand. Samples were loaded in a 10 µL capillary and submitted to a temperature gradient starting at 35°C and finishing at 95°C. The ratio of emission intensities at 350 nm and 330 nm was measured, the resulting melting curves were generated by plotting the first derivative of this ratio against temperature, and the temperature of inflection (Ti) was defined as the maximum of this derivative curve.

### Steady-state fluorescence anisotropy

Fluorescent anisotropy assays were performed using a Safire microplate reader (TECAN) with the excitation wavelength set at 470 nm and emission measured at 530 nm for FITC-labeled peptides. The buffer solution for the assays was 20 mM Tris-HCl pH 7.5, 150 mM NaCl, 5% (v/v) glycerol, and 2 mM DTT. The measurements were initiated at a high protein concentration (10 µM of *Bma*DAF-12 or *Hco*DAF-12), and the protein sample was then repeatedly diluted 2-fold with buffer solution. For each point of the titration curve, the protein sample was mixed with fluorescent peptide (5 nM final concentration) and 20 µM of ligand (two molar excess relative to the highest concentration of protein) for ligand-bound measurements. The reported data are the average of at least three independent experiments, and error bars correspond to standard deviations. Kd values were deduced by non-linear sigmoidal analysis using GraphPad (Prism).

### Cell-based transactivation assays

NIH3T3 cells (ATCC) were cultured in DMEM (Dulbecco’s modified Eagle’s medium) with L-glutamine supplemented with 10% (v/v) FBS containing 100 U/mL penicillin and 100 μg/mL streptomycin. To perform the cell-based transactivation assays, 12.5 × 10^3^ NIH3T3 cells were seeded in white 96-well plates with transparent bottom. After 24 hours, the cells were transiently transfected in 125μL serum-free DMEM with 50 ng of pFN26A_DAF-12-LBD constructs bearing a Renilla luciferase gene used for normalization and 50 ng of pGL4.35 plasmid bearing the luciferase reporter gene under the control of the Gal4 response element (UAS, upstream activation sequence) (Promega) using 0.3 μL of TransIT-X2 transfection reagent (Mirus Bio) in 10μL of Opti-MEM. Serum-free medium was replaced 5 hours later by 200 μL of complete medium with ligands or vehicle control. The final concentration of DMSO was maintained at 0.1% in each well. After an incubation of 24 hours, the cells were lysed and luciferase and renilla activities were successively measured using the Dual-Glo luciferase assay system (Promega) with a FLUOstar OMEGA microplate reader (BMG Labtech). In order to draw a dose response curve, we first calculated the ratio of luminescence from the luciferase reporter to luminescence from the renilla reporter for each condition and replicate. The ratio of DAF-12-transfected cells was normalized to the ratio of empty vector-transfected cells that were treated under the same conditions. The curve was fitted using GraphPad Prism 8 under a non-linear regression fit based on a sigmoidal dose-response with variable slope.

### Crystallization and structure resolution

Crystals of *Bma*DAF-12 and *Hco*DAF-12 complexes with Δ4-DA and peptides derived from coactivators (PGC1ɑ-1 and SRC2-2, respectively) were obtained by co-crystallization. 100 nL of protein concentrated at 13.5 mg/mL and 9.6 mg/mL with two molar equivalents of Δ4-DA and two molar equivalents of PGC1ɑ-1 for *Bma*DAF-12 and with three molar equivalents of Δ4-DA and five molar equivalents of SRC2-2 for *Hco*DAF-12 were mixed with 100 nL of precipitant solution (0.2 M ammonium citrate dibasic, 20% (w/v) PEG 3350 for *Bma*DAF-12 and 2.4 M sodium malonate pH 7 for *Hco*DAF-12) and equilibrated against a reservoir of 50 µL of precipitant solution, using the sitting-drop vapor diffusion technique. Crystals appeared in 24 hours. Crystals were mounted from mother liquor onto a cryoloop, soaked in the precipitant solution containing 20% (v/v) glycerol and flash-frozen in liquid nitrogen. Diffraction data were collected at the ID-30B beamline at the European Synchrotron Radiation Facilities (λ = 0.969 Å, 100 K) at 1.70 Å resolution for *Bma*DAF-12 and 2.0 Å resolution for *Hco*DAF-12. Diffraction data were automatically processed using the Information System for Protein Crystallography Beamlines (ISPyB). The initial phases were obtained by automatic molecular replacement using Phenix.phaser [24] with the Alphafold [20] model of the corresponding complexes. Models were built with Coot [25] and refined with Phenix.refine [24]. Figures were prepared with PyMOL (The PyMOL Molecular Graphics System, Version 3.0 Schrödinger, LLC.).

### Phylogenetic analysis and sequence logos

Full-length amino acid sequences of DAF-12 were recovered from WormBase ParaSite GeneTree WBGT00950000409411. After initial curation, partial sequences (lacking clearly-defined DBD and/or LBD domains) were removed from the analysis. Accession numbers of the retained sequences are provided in S5 Table. An initial amino acid alignment was performed using Clustal Omega [26] (S1 File), which was followed by automated refinement using BMGE [27] (S2 File). The phylogenetic tree was calculated based on the final alignment of 118 sequences using 331 amino acid positions and allowing gaps. Phylogenetic relationships were assessed using the Maximum Likelihood (ML) method as implemented in RAxML [27]. The ML tree was calculated by applying a LG matrix [28], allowing for a proportion of invariable sites (+I) and a discrete Gamma distribution (+G4). The robustness of each node of the resulting tree was assessed by rapid bootstrap analyses (with 1,000 pseudoreplicates). The tree was visualized using iTOL [28] rooted with the *Plectus sambesii* DAF-12 sequence. The image output of the alignment was generated using Unipro UGENE [28], with a Clustal X coloring palette and a sequence order based on the results of the phylogenetic tree. Sequence logos for the sequence stretches corresponding to the LBD of *B. malayi* and *H. contortus* were calculated for the entire untrimmed alignment using WebLogo3 [28].

## Acknowledgements

We acknowledge the European Synchrotron Radiation Facility (ESRF) for the provision of synchrotron radiation facilities, and we thank the staff of the ESRF and EMBL Grenoble for assistance and support in using beamline ID30 under Proposal No MX-2600. The Center for Structural Biology is supported by the French Infrastructure for Integrated Structural Biology (FRISBI), a national infrastructure supported by the French National Research Agency (ANR-10-INBS-05). This work used the PIBBS platform supported by the French National Research Agency (ANR-10-INBS-05). Native MS experiments were carried out at the Montpellier Proteomics Platform (PPM, Biocampus Montpellier) supported by the regional funds FEDER/Région Occitanie, MUSE and Labex EpiGenMed.

## Supporting information

**Figure S1. Biophysical characterization of the *Brugia malayi* (*Bma*) and *Haemonchus contortus* (*Hco*) DAF-12 ligand-binding domains (LBDs). A. and B.** Size exclusion chromatography (SEC) chromatograms of *Bma*DAF-12 LBD (A) and *Hco*DAF-12 LBD (B) with respective 10% SDS-PAGE Bis-Tris gel of the protein after the final purification step. **C.** Alignment of *Hco*DAF-12 and *Ancylostoma ceylanicum* (*Ace*) DAF-12 LBD sequences. Secondary structure elements (H, α-helix; B, β-strand) from the crystal structures are indicated. Cysteines that were mutated into serines in both proteins are highlighted in purple. **D.** Overlay of far-UV circular dichroism (CD) spectra of the *Bma*DAF-12 and *Hco*DAF-12 LBDs. Secondary structure elements coming from the deconvolution of these data are indicated (a) and compared to the theoretical values (b). **E.** Mass-weighted dynamic light scattering (DLS) size-distribution profiles for the purified *Bma*DAF-12 and *Hco*DAF-12 LBDs. The deduced hydrodynamic radii are indicated and are in agreement with the size of the monomeric form of the proteins.

**Figure S2. Ligand binding to the *Brugia malayi* (*Bma*) and *Haemonchus contortus* (*Hco*) DAF-12 ligand-binding domains (LBDs). A. and B**. Nano-DSF analysis of ligand binding to the *Bma*DAF-12 (A) and *Hco*DAF-12 (B) LBDs. Δ4- and Δ7-dafachronic acids (Δ4-DA and Δ7-DA) or DMSO (apo condition) were added in three molar excess to the protein. The first derivative of the ratio between intrinsic fluorescence at 350 nm and 300 nm is plotted against the temperature. The experiments were repeated three times independently. **C. and D.** Native mass spectra of apo *Bma*DAF-12 LBD (C) sprayed at 20 µM, showing one species corresponding to the protein at 28 755 Da (grey circles), and *Bma*DAF-12 in the presence of 50 µM Δ4-DA (D), showing an additional species at 29 170 Da corresponding to one Δ4-DA bound to *Bma*DAF-12 (grey circle with red star).

**Figure S3. Titration curves of affinity measurements on the *Brugia malayi* (*Bma*) and *Haemonchus contortus* (*Hco*) DAF-12 ligand-binding domains (LBDs) by fluorescence anisotropy. A.** Affinity curves for the *Bma*DAF-12 (left) and *Hco*DAF-12 (right) LBDs and human coactivator-derived peptides (Table 1) in the absence of ligand. **B.** Affinity curves for the *Bma*DAF-12 (left) and *Hco*DAF-12 (right) LBDs and human coactivator-derived peptides (Table 1) in the presence of two molar excess of Δ4-DA. **C.** Affinity curves for the *Bma*DAF-12 (left) and *Hco*DAF-12 (right) LBDs and DIP-1-derived peptide in the absence of ligand and in the presence of two molar excess of Δ4-DA and Δ7-DA. **D.** Affinity curves for WT and mutant *Bma*DAF-12 LBDs and PGC1α-1 (left) and DIP-1 (right) peptides in the presence of two molar excess of Δ4-DA. **E.** Affinity curves for *Hco*DAF-12 and *Hco*DAF-12ΔH12 LBDs and SRC2-2 (left) and DIP-1 (right) peptides in the presence of two molar excess of Δ4-DA. **F.** Affinity curves for the *Bma*DAF-12 (left) and *Hco*DAF-12 (right) LBDs and WT and mutant SRC1-1 peptides in the presence of two molar excess of Δ4-DA.

**Figure S4. Conservation of residues involved in coactivator binding in the ligand-binding domains (LBDs) of the human nuclear receptors VDR, LXR, and FXR and DAF-12. A.** Close-up view of the *h*FXR LBD-SRC1 coactivator peptide (CoA) interface from the crystal structure (PDB 3BEJ) [29]. Residues involved in binding are shown as sticks and labeled. **B.** Sequence alignment of the LBDs of the *Brugia malayi* (*Bma*) DAF-12, the *Haemonchus contortus* (*Hco*) DAF-12, the closest human nuclear receptors hVDR, hLXRβ and hFXR and the more distant nuclear receptor, RXRα. Residues involved in coactivator recruitment are highlighted in purple. The two phenylalanines conserved in DAF-12 but not present in human nuclear receptors are framed in red. **C.** Titration curves of affinity measurements on the mammalian LXRβ and RXRα LBDs and WT and mutant SRC1-1 peptides in the presence of two molar excess of agonist ligands (epoxycholesterol and CD3254, respectively), by fluorescence anisotropy.

**Figure S5. Structural conservation of the DAF-12 ligand-binding domains (LBDs) from different nematodes. A.** Superimposition of the *Brugia malayi* (*Bma*) DAF-12-Δ4-DA-PGC1α-1 (PDB 9TLX, pink) structure with the *Haemonchus contortus* (*Hco*) DAF-12-Δ4-DA-SRC2-2 (PDB 9TL4, wheat), the *Strongyloides stercoralis* (*Sst*) DAF-12-Δ4-DA-SRC1-4 (PDB 3GYT, orange), and the *Ancylostoma ceylanicum* (*Ace*) DAF-12-Δ7-DA-SRC2-3 (PDB 3UP0, green) structures. **B.** Table of calculated root mean square deviation (RMSD) over the Cα between the *Bma*DAF-12-Δ4-DA-PGC1α-1, the *Hco*DAF-12-Δ4-DA-SRC2-2, the *Sst*DAF-12-Δ4-DA-SRC1-4, the *Sst*DAF-12-Δ7-DA-SRC1-4, the *Ace*DAF-12-CA-SRC2-3, and the *Ace*DAF-12-Δ7-DA-SRC2-3 structures.

**Figure S6. Close-up view of the *Haemonchus contortus* (*Hco*) DAF-12-DIP-1 interface** from an AlphaFold model of the complex generated using the Alphafold3 server [20]. Residues involved in the interaction are shown as sticks and labeled.

**Figure S7. Phylogenetic tree of DAF-12 proteins and sequence alignment of the ligand binding domain (LBD)** (positions 1880 to 2207 of the untrimmed alignment).

**Figure S8. Clade-specific presentation of the sequence logos for the DAF-12 ligand binding domain (LBD)** (positions 1897 to 2198 of the untrimmed alignment).

**Table S1. Thermal stabilization of the *Brugia malayi* (*Bma*) and *Haemonchus contortus* (*Hco*) DAF-12 ligand-binding domains (LBDs) following ligand incubation.** Inflection temperatures deduced from nano-DSF experiments (Figs 1B and S2A and B Fig) of the *Bma*DAF-12 and *Hco*DAF-12 LBDs in their apo state and in the presence of one and three molar excess of Δ4-DA and Δ7-DA.

**Table S2. X-ray data collection and refinement statistics for *Brugia malayi* (*Bma*) and *Haemonchus contortus* (*Hco*) DAF-12.** Data were collected from one crystal for each dataset. Values in parentheses are for the highest-resolution shell.

**Table S3. Ligand binding pocket volumes (Å3) of the ligand-binding domains (LBDs) of human hLXR, hFXR, and hVDR, and of *Brugia malayi* (*Bma*) DAF-12, *Haemonchus contortus* (*Hco*) DAF-12, *Ancylostoma ceylanicum* (*Ace*) DAF-12, and *Strongyloides stercoralis* (*Sst*) DAF-12.** Volumes were estimated using KVFinder-web.

**Table S4. Conservation of hydrophobic amino acids involved in ligand binding in DAF-12 from *Brugia malayi* (*Bma*), *Haemonchus contortus* (*Hco*), *Strongyloides stercoralis* (*Sst*), and *Ancylostoma ceylanicum* (*Ace*).** The last column gives the correspondence with the sequence alignment numbering in Fig 4.

**Table S5. Sequences and accession numbers of the DAF-12 proteins used in this analysis.**

**File S1. Sequence alignment of complete DAF-12 proteins.**

**File S2. Trimmed sequence alignment of DAF-12 proteins used to generate the phylogenetic tree.**

## Notes

### Competing Interest Statement

The authors have declared no competing interest.

## References

1. Hotez PJ, Brindley PJ, Bethony JM, King CH, Pearce EJ, Jacobson J. Helminth infections: The great neglected tropical diseases. Journal of Clinical Investigation. 2008. doi:10.1172/JCI34261

2. Kotze AC, Hunt PW, Skuce P, von Samson-Himmelstjerna G, Martin RJ, Sager H, et al. Recent advances in candidate-gene and whole-genome approaches to the discovery of anthelmintic resistance markers and the description of drug/receptor interactions. Int J Parasitol Drugs Drug Resist. 2014;4: 164–184. doi:10.1016/j.ijpddr.2014.07.007

3. Motola DL, Cummins CL, Rottiers V, Sharma KK, Li T, Li Y, et al. Identification of Ligands for DAF-12 that Govern Dauer Formation and Reproduction in C. elegans. Cell. 2006;124: 1209–1223. doi:10.1016/j.cell.2006.01.037

4. Luciani GM, Magomedova L, Puckrin R, Urbanus ML, Wallace IM, Giaever G, et al. Dafadine inhibits DAF-9 to promote dauer formation and longevity of Caenorhabditis elegans. Nat Chem Biol. 2011;7: 891–893. doi:10.1038/nchembio.698

5. Wang Z, Schaffer NE, Kliewer SA, Mangelsdorf DJ. Nuclear receptors: emerging drug targets for parasitic diseases. J Clin Invest. 2017;127: 1165–1171. doi:10.1172/JCI88890

6. Ma G, Wang T, Korhonen PK, Young ND, Nie S, Ang C-S, et al. Dafachronic acid promotes larval development in Haemonchus contortus by modulating dauer signalling and lipid metabolism. PLOS Pathog. 2019;15: e1007960. doi:10.1371/journal.ppat.1007960

7. Jaleta TG, Lok JB. Advances in the Molecular and Cellular Biology of Strongyloides spp. Curr Trop Med Reports. 2019;6: 161–178. doi:10.1007/s40475-019-00186-x

8. Long T, Alberich M, André F, Menez C, Prichard RK, Lespine A. The development of the dog heartworm is highly sensitive to sterols which activate the orthologue of the nuclear receptor DAF-12. Sci Rep. 2020;10: 11207. doi:10.1038/s41598-020-67466-9

9. Cheong MC, Wang Z, Jaleta TG, Li X, Lok JB, Kliewer SA, et al. Identification of a nuclear receptor/coactivator developmental signaling pathway in the nematode parasite *Strongyloides stercoralis*. Proc Natl Acad Sci. 2021;118. doi:10.1073/pnas.2021864118

10. Wang Z, Zhou XE, Motola DL, Gao X, Suino-Powell K, Conneely A, et al. Identification of the nuclear receptor DAF-12 as a therapeutic target in parasitic nematodes. Proc Natl Acad Sci. 2009;106: 9138–9143. doi:10.1073/pnas.0904064106

11. Lok JB, Kliewer SA, Mangelsdorf DJ. The ‘nuclear option’ revisited: Confirmation of Ss-daf-12 function and therapeutic potential in Strongyloides stercoralis and other parasitic nematode infections. Mol Biochem Parasitol. 2022;250: 111490. doi:10.1016/j.molbiopara.2022.111490

12. Bétous R, Emile A, Che H, Guchen E, Concordet D, Long T, et al. Filarial DAF-12 sense the host serum to resume iL3 development during infection. PLOS Pathog. 2023;19: e1011462. doi:10.1371/journal.ppat.1011462

13. Wang Z, Cheong MC, Tsien J, Deng H, Qin T, Stoltzfus JD, et al. Characterization of the endogenous DAF-12 ligand and its use as an anthelmintic agent in Strongyloides stercoralis. Elife. 2021;10. doi:10.7554/eLife.73535

14. Lonard DM, O’Malley BW. Nuclear Receptor Coregulators: Judges, Juries, and Executioners of Cellular Regulation. Molecular Cell. 2007. doi:10.1016/j.molcel.2007.08.012

15. Ludewig AH, Kober-Eisermann C, Weitzel C, Bethke A, Neubert K, Gerisch B, et al. A novel nuclear receptor/coregulator complex controls *C. elegans* lipid metabolism, larval development, and aging. Genes Dev. 2004;18: 2120–2133. doi:10.1101/gad.312604

16. Zhi X, Zhou XE, Melcher K, Motola DL, Gelmedin V, Hawdon J, et al. Structural Conservation of Ligand Binding Reveals a Bile Acid-like Signaling Pathway in Nematodes. J Biol Chem. 2012;287: 4894–4903. doi:10.1074/jbc.M111.315242

17. le Maire Albane, Bourguet William. Retinoic acid receptors: structural basis for coregulator interaction and exchange. Springer. Retinoic acid Signaling Book. Springer. 2013.

18. Guerra JVS, Ribeiro-Filho H V, Pereira JGC, Lopes-de-Oliveira PS. KVFinder-web: a web-based application for detecting and characterizing biomolecular cavities. Nucleic Acids Res. 2023;51: W289–W297. doi:10.1093/nar/gkad324

19. Goddard TD, Huang CC, Meng EC, Pettersen EF, Couch GS, Morris JH, et al. UCSF ChimeraX: Meeting modern challenges in visualization and analysis. Protein Sci. 2018;27: 14–25. doi:10.1002/pro.3235

20. Abramson J, Adler J, Dunger J, Evans R, Green T, Pritzel A, et al. Accurate structure prediction of biomolecular interactions with AlphaFold 3. Nature. 2024;630: 493–500. doi:10.1038/s41586-024-07487-w

21. Howe KL, Bolt BJ, Shafie M, Kersey P, Berriman M. WormBase ParaSite − a comprehensive resource for helminth genomics. Mol Biochem Parasitol. 2017;215: 2–10. doi:10.1016/j.molbiopara.2016.11.005

22. Gerisch B, Rottiers V, Li D, Motola DL, Cummins CL, Lehrach H, et al. A bile acid-like steroid modulates *Caenorhabditis elegans* lifespan through nuclear receptor signaling. Proc Natl Acad Sci. 2007;104: 5014–5019. doi:10.1073/pnas.0700847104

23. McKenna NJ, Lanz RB, O’Malley BW. Nuclear Receptor Coregulators: Cellular and Molecular Biology*. Endocr Rev. 1999;20: 321–344. doi:10.1210/edrv.20.3.0366

24. Liebschner D, Afonine P V., Baker ML, Bunkóczi G, Chen VB, Croll TI, et al. Macromolecular structure determination using X-rays, neutrons and electrons: recent developments in *Phenix*. Acta Crystallogr Sect D Struct Biol. 2019;75: 861–877. doi:10.1107/S2059798319011471

25. Emsley P, Lohkamp B, Scott WG, Cowtan K. Features and development of Coot. Acta Crystallogr Sect D Biol Crystallogr. 2010;66: 486–501. doi:10.1107/S0907444910007493

26. Sievers F, Wilm A, Dineen D, Gibson TJ, Karplus K, Li W, et al. Fast, scalable generation of high-quality protein multiple sequence alignments using Clustal Omega. Mol Syst Biol. 2011;7. doi:10.1038/msb.2011.75

27. Criscuolo A, Gribaldo S. BMGE (Block Mapping and Gathering with Entropy): a new software for selection of phylogenetic informative regions from multiple sequence alignments. BMC Evol Biol. 2010;10: 210. doi:10.1186/1471-2148-10-210

28. Le SQ, Gascuel O. An improved general amino acid replacement matrix. Mol Biol Evol. 2008;25: 1307–1320. doi:10.1093/molbev/msn067

29. Soisson SM, Parthasarathy G, Adams AD, Sahoo S, Sitlani A, Sparrow C, et al. Identification of a potent synthetic FXR agonist with an unexpected mode of binding and activation. Proc Natl Acad Sci. 2008;105: 5337–5342. doi:10.1073/pnas.0710981105

